# Genetic Determinants of Cell-Type-Specific Responses to Cardiotoxic Chemotherapeutic Agents

**DOI:** 10.64898/2026.10.16.759549

**Authors:** Erik McIntire, Kenneth A. Barr, Natalia M. Gonzales, Olivia L. Allen, Jonathan Burnett, Yoav Gilad

## Abstract

Doxorubicin (**DOX**) and 5-fluorouracil (**5-FU**) are effective for treating a variety of cancers, yet many patients taking these drugs experience adverse cardiovascular events that are severe enough to pause treatment, often prolonging the course of chemotherapy and reducing its efficacy. Individual risk for drug-induced cardiotoxicity is likely mediated, at least in part, by genetic variants that control how the heart and vasculature respond to drug-induced perturbations. To identify loci associated with drug-induced cardiotoxicity, we developed an *in vitro* model that allows us to characterize the effects of chemotherapeutic drugs on a wide range of cardiac cell types, including cardiomyocytes, endothelial cells, and fibroblasts. Using single-cell RNA-seq, we profiled responses to DOX and 5-FU across 46 individuals and 14 cell types, identifying hundreds of expression quantitative trait loci that modify cardiovascular transcription in response to each drug (**response eQTLs**). Drug-responsive eQTL genes include genes that were previously linked to drug-induced cardiotoxicity and cardiovascular disease, such as *ABCB4*, *HAS3*, *TNFRSF1A*, and *OGFR*. Over a third of drug-responsive genes are shared between the DOX and 5-FU treatments, suggesting that genes predictive of DOX-induced cardiotoxicity may extend to 5-FU. Among the credible colocalizations we identified between GWAS signals and drug-response eQTLs, we discovered a locus associated with ischemic heart disease that is also a DOX-responsive eQTL for *IL6R.* IL6R was previously nominated as a druggable target for heart disease, and its binding partner IL6 is a candidate biomarker for anthracycline-induced cardiotoxicity. These findings advance the discovery of genetic variants underlying cardiotoxicity and support development of genotype-based predictors for treating cancer.

## INTRODUCTION

The leading causes of death worldwide are cardiovascular disease and cancer^1^. Patients taking chemotherapeutic drugs often experience side effects that compromise the function of the heart and blood vessels, compounding the risk of cardiovascular disease^2,3^. While widely-prescribed drugs like doxorubicin (**DOX**, an anthracycline) and 5-fluorouracil (**5-FU**, an anti-metabolite) are effective for treating a variety of cancers^4^, as many as 48% of patients on DOX and 35% on 5-FU will experience arrhythmia, left ventricular dysfunction, myocardial ischemia, sudden cardiac death, or other symptoms of cardiotoxicity, some of which may emerge well after the treatment has stopped^5–7^. Essentially, many of the same patients who owe their recovery to drugs like DOX or 5-FU, survive cancer only to be burdened with an increased risk for cardiovascular disease. Thus, drug-induced cardiotoxicity often reduces the very gains in quantity and quality of life that chemotherapy is intended to provide^8^.

Despite decades of research on the toxicities associated with drugs like DOX and 5-FU, clinicians remain limited in their ability to predict who is at risk for severe adverse effects. Risk for drug-induced cardiotoxicity generally increases with dose and treatment length, although in some patients, small or even acute doses of DOX or 5-FU can produce adverse effects^2^. Age, sex, mediastinal radiotherapy, and cardiovascular health may further modify individual responses to cardiotoxic drugs, but these factors cannot be used to forecast specific symptoms, nor can they predict the severity of cardiotoxicity^9,10^. Patients vary widely in their responses to both DOX and 5-FU, it is reasonable to assume that drug-induced cardiotoxicity risk is mediated, at least in part, by genetic factors. Indeed, while the genetic component of 5-FU-induced cardiotoxicity remains understudied^11^, 180 genetic variants have been associated with DOX-induced cardiotoxicity^12^. Additionally, *in vitro* studies of human cardiomyocytes have identified hundreds of expression quantitative trait loci (**eQTLs**) that regulate gene expression levels specifically in response to DOX treatment^13–15^. Because they modulate gene regulation in a drug-dependent manner, drug response QTLs can provide insights into drug mechanisms and potentially serve as functional biomarkers for cardiotoxicity and disease risk^16^.

A growing body of work has shown that cardiomyocytes derived from induced pluripotent stem cells (**iPSCs**) can recapitulate aspects of anthracycline cardiotoxicity *in vivo*^13,14^; indeed, we previously used this system to identify eQTLs that modulate transcriptional responses to DOX^15^. While differentiated cardiomyocytes remain a choice model for studies of drug-induced cardiotoxicity, the diverse phenotypes observed in patients treated with DOX or 5-FU suggest that these drugs might adversely affect multiple cell types. For instance, DOX can produce vascular toxicity by increasing endothelial cell permeability and it can cause cardiac fibrosis by activating cardiac fibroblasts^17^. Similarly, 5-FU has been associated with coronary vasospasm, endothelial dysfunction, and thrombosis^6^, which are likely to involve vascular cell types in addition to cardiac muscle.

With these observations in mind, we wanted to develop an *in vitro* system that would allow us to explore transcriptional responses to DOX and 5-FU across multiple cell types – particularly because many of the eQTLs that regulate these responses are expected to be cell-type-specific^16^. In a recent eQTL mapping study^18^, we used an unguided differentiation protocol^19^ to generate heterogeneous differentiating cultures (**HDCs**) comprising dozens of cell types from each germ layer. By combining this protocol with methods for generating cardiac organoids, we developed a way to enrich HDCs for cardiovascular cell types. Using ‘cardiac HDCs’, as we refer to this new model, we were able to measure transcriptional responses to DOX and 5-FU in cell types other than cardiomyocytes, revealing a complex regulatory network that affects multiple aspects of cardiovascular function.

## RESULTS

### Cardiac HDCs are highly diverse and enriched for cardiovascular cell types

To measure the transcriptional effects of DOX and 5-FU across a range of cell types and individuals, we used a panel of 49 human iPSCs^20^ to generate cardiac HDCs, which we cultured for 14 days (**Figure 1**). On day 15, we independently treated each cell line with either 1 µM DOX, 5 µM 5-FU, or a vehicle control (dimethyl sulfoxide, i.e., DMSO) for 24 hours. On day 16, we dissociated the cells and prepared them for single-cell RNA-sequencing (**scRNA-seq**) on the 10x Genomics platform (**Figure 1A**, **Methods**). After filtering and quality control, we retained 7.94 billion unique molecular identifier (**UMI**) counts from 46 individuals across all three treatment groups (an average of 12,117 UMI counts per cell). In total, we retained high-quality data from 668,281 cells, including 208,092 DOX-treated cells, 242,913 5-FU-treated cells, and 217,276 control cells.

**Figure 1.**
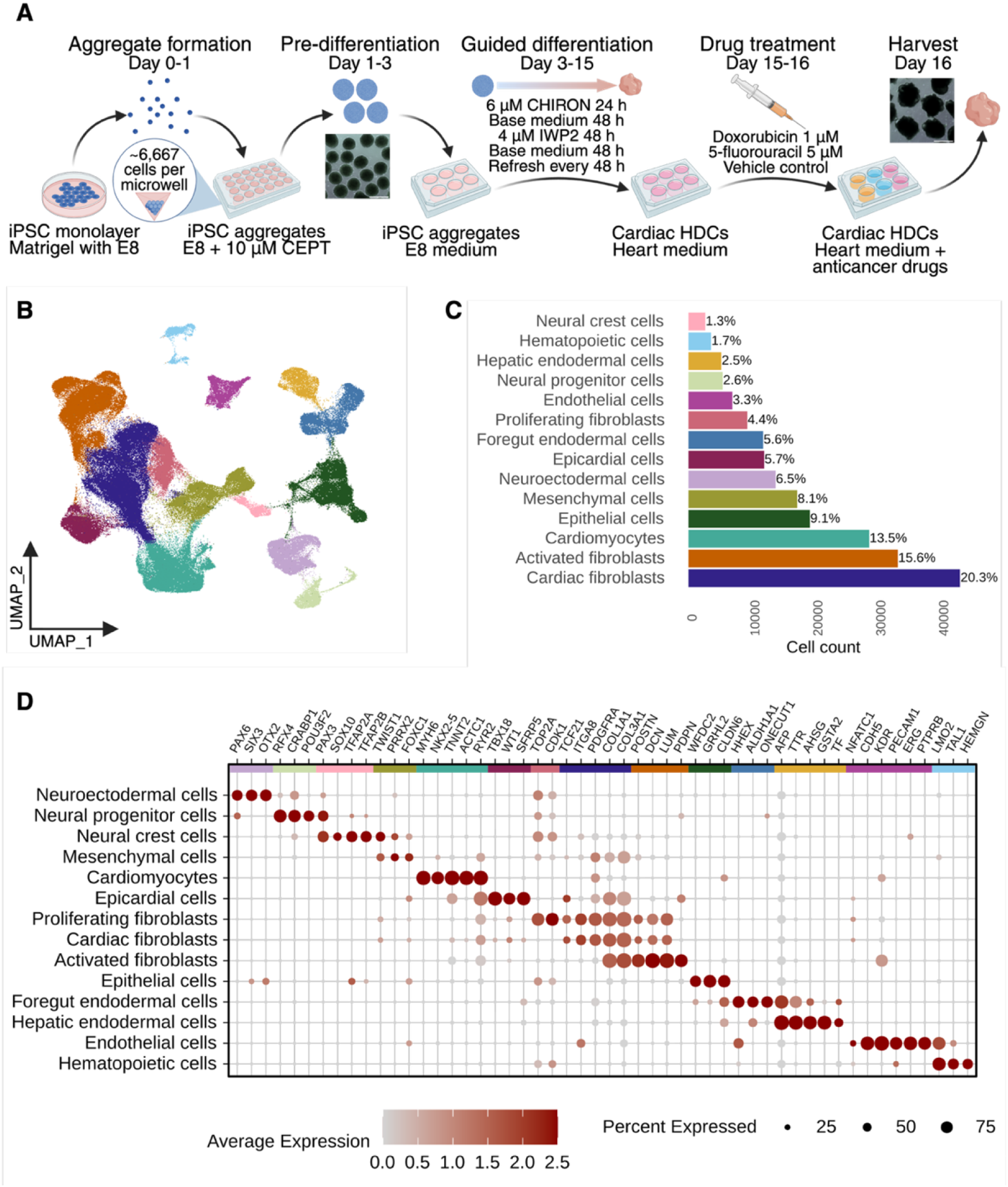
Guided differentiation of iPSCs generates cardiac HDCs harboring diverse cell types. (**A**) Guided differentiation and treatment of cardiac HDCs. To generate cardiac HDCs, we replace the typical ROCK inhibitor with the cytoprotectant CEPT to form three-dimensional iPSC aggregates measuring 300 μm in diameter. CEPT is critical for cell viability and maturation in addition to producing consistent cell type proportions between batches and individuals. To promote recovery after formation, we move aggregates to suspension culture in E8 medium. For 48 h, we avoid replenishing media to facilitate a reduction in thermolabile FGF activity, thereby easing cells out of pluripotency and poising them for differentiation. Temporally modulated WNT signaling with CHIR99021 and IWP2 is used as the differentiation stimulus. The omission of insulin is key to cardiac HDC diversity, as it facilitates nonmyocyte differentiation. On day 15 of culture, we treated cardiac HDCs with either 1μM DOX, 5μM 5-FU, or vehicle control (DMSO) for 24 h before harvest for scRNA-seq. Scale bars: 500μm. Abbreviations: E8: Essential 8 medium. CEPT: Chroman 1, emricasan, polyamines, and trans-ISRIB. Base heart medium: RPMI 1640 Medium, GlutaMAX Supplement, HEPES, 2% v/v B-27 Supplement, minus insulin. FGF: fibroblast growth factor. DMSO: dimethyl sulfoxide. (**B**) UMAP plot of 217,276 sequenced cells from 46 iPSC lines in the control data, manually annotated. (**C**) Proportions of the 14 cell types identified in the control data. (**D**) Dot plot showing relative expression of key marker genes for each cell type in the control data.

To compare gene expression levels across different cell types and treatments, we needed an annotation strategy capable of producing labels that are robust to the transcriptional changes introduced by DOX and 5-FU (e.g., a successful approach would correctly identify a DOX-treated cardiomyocyte as a cardiomyocyte rather than classifying it as a different cell type). To begin, we log-normalized gene expression values from each treatment dataset separately. In the control data, we performed scaling and regressed out cell cycle phase differences. This step minimizes variability among proliferating cells while preserving phase differences between cycling and non-cycling cells, thereby maintaining distinctions between progenitor and terminally differentiated populations (**Methods**). Principal components analysis (**PCA**) and unsupervised clustering yielded 14 cell clusters (**Figure 1B-C**), which we annotated manually according to the expression of marker genes (**Figure 1D**, **Methods**). Following this manual annotation, we applied Seurat^21^ to predict cell type labels in the DOX and 5-FU datasets, leveraging the annotated control cells as a reference dataset.

Using this approach, we annotated 14 cell types representing all three germ layers in cardiac HDCs, including cardiomyocytes, neural crest cells, and foregut endodermal cells (**Figure 1B-D**), each of which were represented in all of the individuals we sampled (**Supplementary Figure 1**). Marker gene expression patterns are consistent among matched cell types from the control and drug treatment groups (**Supplementary Figure 2A**), and both DOX-treated and 5-FU-treated cells show high-confidence cell type prediction score distributions (**Supplementary Figure 2B-C**). Moreover, cardiac HDCs from the DOX and 5-FU groups exhibit similar cell type diversity across individuals (**Supplementary Figures 3-4**), indicating that our annotation strategy is robust.

### Cardiac HDC responses to DOX and 5-FU are consistent with known mechanisms of cardiotoxicity

Before exploring genetic effects on gene regulation, we compared the gene expression profiles of drug-treated cells to control-treated cells and identified, for each cell type, genes that are differentially expressed (**DE**) between either DOX or 5-FU treatments and the control group. Throughout the results, we use the term ‘condition’ to describe cells belonging to a particular cell type and treatment group; for example, control-treated epicardial cells and 5-FU-treated epicardial cells are distinct conditions; DOX treated cardiomyocytes and DOX treated epicardial cells are also distinct conditions.

DOX treatment induced widespread transcriptional changes across all cell types, with the greatest number of DE genes found in activated fibroblasts, cardiac fibroblasts, and cardiomyocytes (**Supplementary Figure 5A**). Treatment with 5-FU produced subtler effects on gene regulation compared to those observed for DOX. In the 5-FU group, the greatest number of DE genes were found in neuroectodermal cells and endothelial cells (**Supplementary Figure 5B**). To broadly characterize the transcriptomic effects of DOX and 5-FU, we used Fast Gene Set Enrichment Analysis (**fgsea**)^22^ to test DE genes from each drug-treated cell type for enrichment among the MSigDB Hallmark pathways^23^ (**Methods**). As expected, we found signatures of cell death and senescence to be prevalent among DE genes from all drug treatment conditions, although some cell types exhibit dissimilar transcriptional responses to DOX and 5-FU (**Figure 2A, Supplementary Tables 1-2**). Across a majority of cell types from both drug treatment groups, DE genes are consistently enriched for the P53-related pathway, depleted of E2F targets, and depleted of G2M checkpoint genes (p.adj < 0.05; **Figure 2A**, **Supplementary Tables 1-2**). However, in the 5-FU treatment group, DE genes from epicardial, mesenchymal, and foregut endoderm cells deviate from this pattern, showing a strong enrichment for E2F targets (epicardial NES=3.052, p.adj=1.74×10^-4^; mesenchymal NES=2.010, p.adj=8.88×10^-5^; foregut endoderm NES=2.362, p.adj=1.33×10^-4^) and G2M checkpoint genes (epicardial NES=2.873, p.adj=1.74×10-4; mesenchymal NES=1.721, p.adj=4.01×10^-4^; foregut endoderm NES=2.256, p.adj=1.33×10^-4^; **Figure 2B-C**). The DE genes in these conditions are similarly depleted in several other pathways, including TNFA signaling via NFKB, epithelial-mesenchymal transition, hypoxia, apoptosis, coagulation, IL6 JAK STAT3 signaling, IL2 STAT5 signaling, and inflammatory response pathways (**Figure 2A, Supplementary Tables 1-2**).

**Figure 2.**
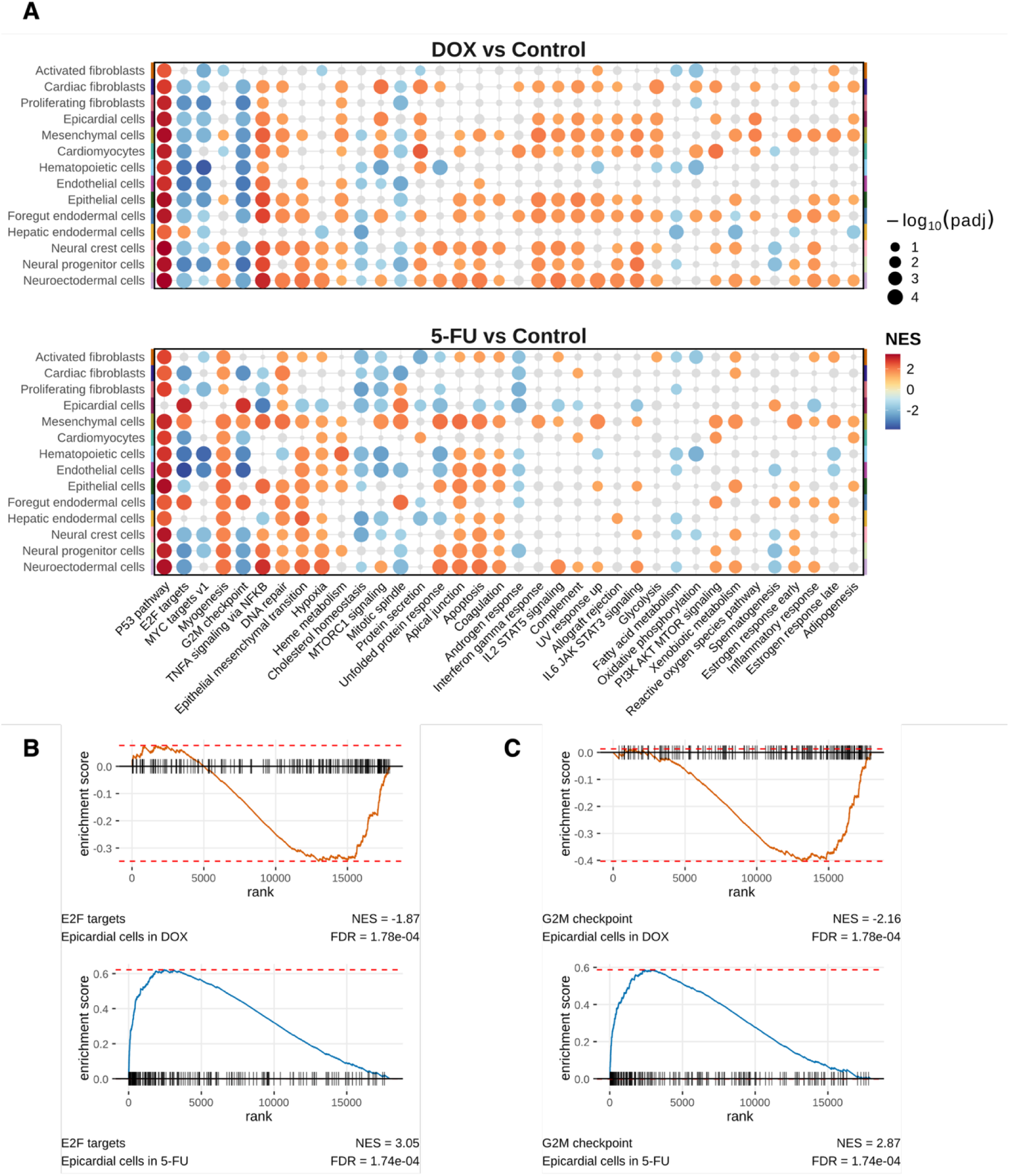
DOX and 5-FU induce widespread signatures of cardiotoxicity, with 5-FU triggering a distinct cell cycle response in epicardial cells. (**A**) Hallmark pathway gene set enrichment analysis results for DE genes identified in DOX-treated vs control-treated cells (top) and 5-FU-treated vs control-treated cells (bottom). Colored circles represent significantly enriched pathways (FDR<0.05) in each cell type, with circle size corresponding to significance level and color corresponding to Normalized Enrichment Score (NES); grey circles represent non-significant pathways. Pathways with <6 significantly perturbed cell types in a treatment group are not shown. (**B**) DE genes from DOX-treated endocardial cells are depleted of E2F target pathway genes (top, orange), while DE genes from 5-FU-treated endocardial cells are enriched for E2F target genes (bottom, blue). (**C**) DE genes from DOX-treated endocardial cells are depleted of G2M checkpoint genes (top, orange), while DE genes from 5-FU-treated endocardial cells are enriched for G2M checkpoint genes (bottom, blue).

In some conditions, DE genes exhibit more specific enrichment patterns that are consistent with known mechanisms of drug cardiotoxicity. For example, many DOX-treated cell types harbor DE genes that are enriched for pathways related to iron handling and inflammation^24^ (i.e., heme metabolism, interferon gamma response, IL6 JAK STAT3 signaling, TNFA signaling via NFKB, PI3K AKT MTOR signaling, and inflammatory response) (**Figure 2A, Supplementary Table 1**). In particular, DE genes identified in DOX-treated cardiomyocytes are enriched for pathways typical of anthracycline-induced cardiotoxicity^2,24^, such as oxidative phosphorylation (NES=1.596, p.adj=0.003), PI3K AKT MTOR signaling (NES=2.100, p.adj=1.98×10^-4^), and reactive oxygen species (NES=1.562, p.adj=0.039). Conversely, DE genes in 5-FU-treated cell types tend to be enriched for remodeling-related pathways (i.e., myogenesis, epithelial mesenchymal transition, apical junction) and pathways indicative of ischemic injury, such as hypoxia and coagulation^6^ (**Figure 2A, Supplementary Table 2**). Together these observations suggest that treatment with an acute dose of DOX or 5-FU is sufficient to produce *in vitro* transcriptional responses that are consistent with the reported molecular effects of each drug.

### Genetic regulatory effects are broadly shared across cell types and treatments

In at-risk individuals, the *in vivo* side effects induced by treatment with DOX or 5-FU can take many forms, such as cardiac arrest, arrythmia, or thrombosis^2^. This heterogeneity led us to suspect that the genetic variants underlying inter-individual differences in cardiotoxicity risk are likely to have cell-type-specific effects on gene expression. To identify regulatory loci with effects that are specific to both treatment and cell type, we separately mapped *cis*-eQTLs for each of the conditions present in our data. First, we filtered the data to ensure sufficient power to detect *cis*-eQTLs, excluding individuals that contributed fewer than 15 cells to a given treatment × cell type condition. We also required that each condition be represented by at least 25 individuals. Four cell types failed to meet these criteria and we excluded the corresponding data from further analyses: hematopoietic cells, neural crest cells, neural progenitors, and neuroectodermal cells (**Methods**). After filtering and re-normalizing the remaining data, we used linear models implemented in MatrixEQTL^25^ to map *cis*-eQTLs (MAF >5%) for each condition, testing variants within 250 kb windows upstream and downstream of each gene’s transcription start site (**TSS**). In addition to sex, we included between two and 12 gene expression principal components as model covariates, which we selected based on the maximum number of *cis-*eQTL genes (**eGenes**) discovered using each model (**Supplementary Figure 6, Methods**). The full set of MatrixEQTL results are provided in **Supplementary File 1.**

Following initial *cis*-eQTL mapping with MatrixEQTL, we implemented multivariate adaptive shrinkage (mash)^26^ to share information across conditions, stabilize per-condition effect size estimates, and detect reproducible between-condition differences while accounting for uncertainty in the estimated effects. Of 12,927 genes tested, we identified 3,014 unique eGenes associated with at least one significant *cis*-eQTL (lfsr <0.1) across the ten cell types and three treatments (**Supplementary Table 3**). The number of eGenes varies across conditions, ranging from 1,373 in 5-FU-treated epithelial cells to 1,657 in DOX-treated cardiomyocytes. In general, the number of eGenes we identified is positively correlated with the number of individuals (R=0.54, P=0.002) (**Figure 3A**) and median cell count (R=0.47, P=0.008) (**Figure 3B**), suggesting that the number of eQTLs identified for each condition can be explained, in part, by differential power. Nonetheless, we observed that for all three treatments, the number of eGenes identified in cardiomyocytes surpassed that of both activated fibroblasts and cardiac fibroblasts, the two most abundant cell types in the data (**Figure 1C**).

**Figure 3.**
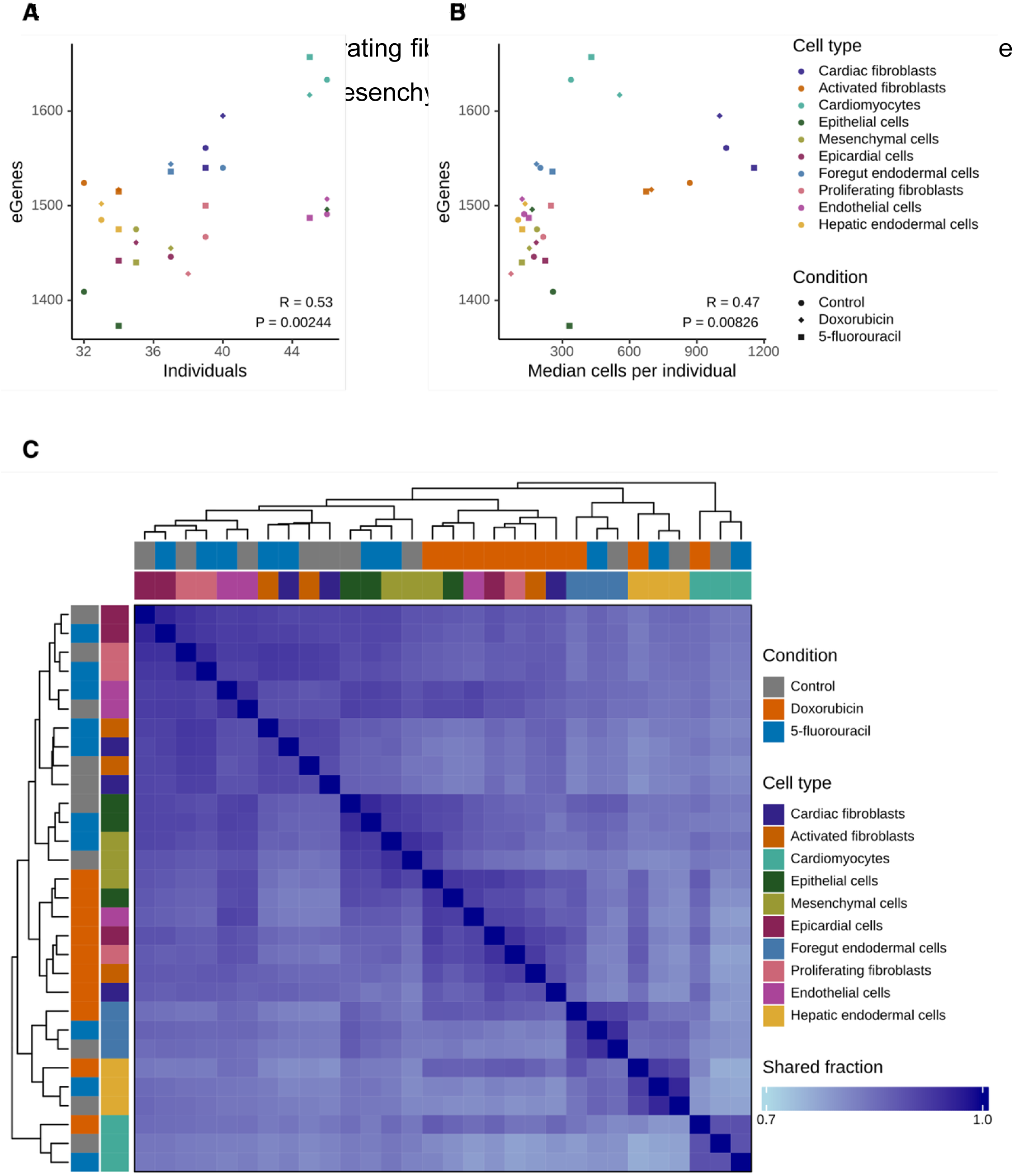
eGenes and corresponding regulatory effects are broadly shared across cell types and treatments. The number of eGene discoveries shows a statistically significant, moderate correlation with the number of individuals (**A**) and median cell number (**B**). (**C**) For each pair of cell-type-treatment conditions, we plotted the percentage of shared eQTLs using the unique eQTLs that met the significance threshold (lfsr <0.1). eQTLs qualify as shared if they are significant in at least one of two conditions, share the same effect sign, and if effect sizes are within a factor of 2.

Next, we investigated the extent to which regulatory effects are shared across the 30 conditions we examined (i.e., spanning control and two treatments in ten cell types). For each pairwise comparison, we assessed only the eQTLs that were significant in at least one of the two conditions. We considered an eQTL to be shared if it had the same effect sign and a similar effect size (i.e., within a factor of 2). Consistent with previous studies that used a similar approach^26,27^, we observed that a high proportion of gene regulatory effects are shared across conditions (**Figure 3C**), with pairwise sharing ranging from 75.7% to 95.8% (median of 85.6%). We found that DOX treatment produces gene regulatory effects that are shared across a majority of cell types, distinguishing them from matched cell types in the control and 5-FU groups (**Figure 3D**). However, for endodermal cells and cardiomyocytes, the effect of cell type on sharing is stronger than the effect of DOX treatment, which is broadly the case for both the 5-FU and control groups. We also observed similar patterns of eQTL sharing among closely-related cell types, including epicardial cells and proliferating fibroblasts, activated and cardiac fibroblasts, and the less mature populations of epithelial and mesenchymal cells (**Figure 3D**).

### Drug-response eGenes are largely cell-type-specific and include known cardiac dysfunction risk genes

Regulatory effects that modify gene expression specifically in response to treatment with DOX or 5-FU may reveal genes, pathways, and cell types that are relevant to the drug’s cardiotoxic side effects. Due to our small sample size, we used a marginal eQTL test in place of an explicit genotype × treatment interaction model to identify response eQTLs, as we and others have done previously^28–31^ (**Methods**). Briefly, this approach leverages the posterior mean effect estimates output by mash, which employs a Bayesian framework to shrink the raw, unstable regression coefficients output by MatrixEQTL, retaining only the strongest signals^26^. The resulting posterior mean effects are effectively de-noised, allowing us to use the log-fold change in eQTL effects between conditions to distinguish between response and non-response eQTLs. Specifically, we compared marginal eQTL effect sizes between matched cell types from the control group and either of the two drug treatment groups. We classified an eQTL as responsive if it was significant (lfsr <0.1) in either the control or the drug group, and if the magnitude of its effect size differed by a fold change >4. We also classified eQTLs as responsive if they were significant in both treatment groups but had differently signed effects.

Recall that we classified eQTLs as shared between conditions when the effect size between conditions differed by less than 2. The choice of the effect size cutoffs to classify shared and response eQTLs are arbitrary. The choice to present results by classifying response eQTLs with the specific effect size cutoff of 4 was made such that shared eQTLs are maximally enriched near TSSs and response eQTLs are distributed equally across the 500 kb region, a pattern that has been observed previously^32^ (**Methods**). That said, this enrichment pattern as well as all subsequent results are robust with respect to a range of effect size cutoffs (between 2 to 10-fold; **Supplementary Figures 6-7**).

After classifying response eQTLs independently for DOX and 5-FU, we identified a total of 912 unique response eQTL eGenes, including 665 unique response eGenes for DOX (**Supplementary Table 4**) and 573 for 5-FU (**Supplementary Table 5**). As expected, response eQTLs exhibit distinct genetic architectures compared to non-response eQTLs^32^. Relative to non-QTL SNPs, both response eQTLs and non-response eQTLs are enriched among candidate *cis* regulatory elements, with response eQTLs showing greater enrichment in enhancer-like regions and non-response eQTLs showing greater enrichment in promoter regions (**Supplementary Table 6**, **Supplementary Figure 8**). DOX response eQTLs show greater enrichment in enhancer-like signatures (OR=1.58, P=6.68×10^−7^) than DOX non-response eQTLs (OR=1.37, P=5.80×10^−9^), and 5-FU response eQTLs display a similar pattern (5-FU response eQTLs: OR=1.65, P = 3.65×10^−7^; 5-FU non-response eQTLs: OR=1.35, P=5.59×10^−9^). When decomposing enhancer elements into proximal versus distal categories, response eQTLs maintain greater enrichment relative to non-response eQTLs for both DOX and 5-FU treatments (**Supplementary Tables 4-5**). Conversely, in promoter-like signatures, non-response eQTLs show greater enrichment than response eQTLs for both DOX (DOX non-response eQTLs: OR=6.04, P=6.45×10^−33^; DOX response eQTLs: OR=4.42, P=3.82×10^−7^) and 5-FU (5-FU non-response eQTLs: OR=6.18, P=4.49×10^−36^; 5-FU response eQTLs: OR=3.17, P=0.001). Given these differences in regulatory architecture, we predicted that response eQTLs would tend to have cell-type-specific effects; accordingly, we observed that within each treatment group, relatively few response eGenes are shared across cell types (**Figure 4A-B**). This is unlikely to be due to power, because we observed that non-response eQTLs are much more often shared among cell types.

**Figure 4.**
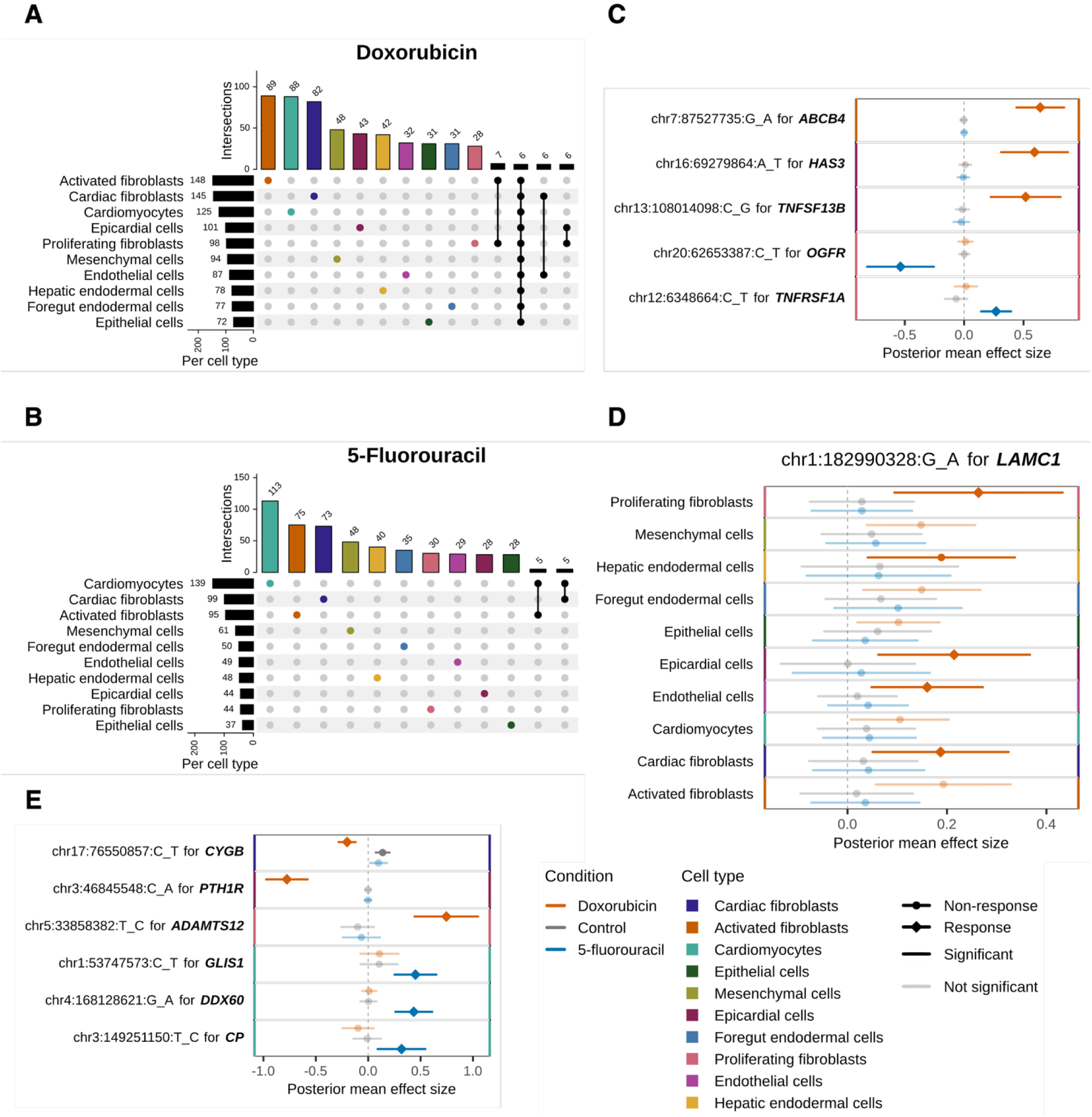
Response eGenes predominantly correspond to distinct cell type lineages and include genes previously implicated in cardiovascular function. Upset plots showing the sharing patterns of drug-response eGenes for (**A**) DOX and (**B**) and 5-FU. Number of response eGenes per cell type are shown on the left of each Upset plot, with intersections on top. Intersections of <5 response eGenes are not shown. (**C**) Response eQTL effect sizes are plotted for five eGenes related to drug-induced cardiotoxicity, three of which are specific to DOX (orange) and two of which are specific to 5-FU. (**D**) A response eQTL specific to DOX has a visible effect on the expression of the cardiotoxicity-associated gene *LAMC1* across all ten cell types. (**E**) Response eQTL effect sizes for three DOX-responsive eGenes and three 5-FU-responsive eGenes that were previously linked to cardiovascular disease phenotypes. For each condition, significant effects (lfsr <0.1) are shown as solid bars and non-significant effects are semi-transparent bars. A diamond shape indicates a drug-responsive eQTL for a given cell-type-treatment condition while a circle indicates a non-responsive eQTL.

A subset of response eQTLs have effects that are present in a single drug treatment context, several of which regulate eGenes that were previously implicated in anthracycline-induced cardiotoxicity risk (**Figure 4C**). For example, we identified *ABCB4* as a DOX-response eGene specific to activated fibroblasts. *ABCB4* encodes a protein that transports DOX and is a known risk gene for anthracycline-induced cardiotoxicity^33^. We also classified the extracellular matrix protein *HAS3* as a DOX response eGene specific to epicardial cells. *HAS3* expression was inversely correlated with cardiotoxicity risk in a study of anthracycline-treated patients^34^, and both *HAS3* and *ABCB4* were functionally validated as DOX-induced cardiotoxicity genes in knockout studies^12^. Additionally, we identified the DOX-response eGene *TNFSF13B* (an immune signaling ligand) in epicardial cells and *LAMC1* (an extracellular matrix component) as a DOX-response eGene in proliferating fibroblasts, epicardial cells, endothelial cells, and cardiac fibroblasts (**Figure 4D**). Both TNFSF13B and LAMC1 were previously nominated as protein biomarkers for anthracycline-induced changes in left ventricular ejection fraction, a hallmark of cardiotoxicity^35,36^. These findings confirm that DOX alters gene expression levels across a variety of cell types. Many of the affected genes are likely to exhibit cell-type-dependent responses that have not been reported in the cardiotoxicity literature, as the majority of *in vitro* studies have focused solely on cardiomyocytes.

While DOX-induced cardiotoxicity has been studied extensively in both humans and animals, relatively little is known about the genes that mediate 5-FU-induced cardiotoxicity^11^. Our ability to identify previously known cardiotoxicity-relevant 5-FU response eGenes was therefore limited by the sparsity of existing data on 5-FU. Nevertheless, we identified two 5-FU-specific response eGenes associated with drug-induced cardiotoxicity in general (**Figure 4C**). The opioid growth factor receptor *OGFR* and tumor necrosis factor receptor *TNFRSF1A* were demonstrated to drive DOX-induced cardiotoxicity in mice and iPSC-derived cardiomyocytes, respectively^37,38^.

We also identified novel response eGenes that lack an established role in drug-induced cardiotoxicity but do link to cardiovascular disease more broadly (**Figure 4E**). For example, among DOX response eGenes we identified the globin gene *CYGB*, which regulates blood pressure and vascular tone^39^; the peptide receptor *PTH1R*, implicated in pulmonary hypertension^40^; and the metalloprotease *ADAMTS12*, a candidate gene for pediatric stroke^1^. We also discovered several 5-FU-specific response eGenes in cardiomyocytes that are relevant to cardiovascular disease: the zinc finger protein *GLIS1* is a risk gene for heart valve disease^41^, the RNA helicase *DDX60* plays a cardioprotective role following myocardial infarction^42^, and serum levels of the copper-binding ferroxidase *CP* predict adverse cardiac events in patients^43^.

While most response eGenes are cell-type-specific, they are markedly less treatment-specific: roughly 36% (n=326) of all response eGenes are shared between the DOX and 5-FU groups (**Figure 5A**). We examined each cell type separately to identify response eGenes that are shared by both the DOX and 5-FU groups, finding several genes that were previously implicated in drug-induced cardiotoxicity (**Figure 5B**). For instance, in cardiomyocytes the shared response eGene *FDXR* was associated with anthracycline-induced cardiotoxicity in pediatric patients, and reduced *FDXR* expression caused mitochondrial iron overload in a separate knockdown study^44,45^. In endothelial cells, the shared response eGenes *ZNF521* and *EYS* have each been linked to anthracycline-and trastuzumab-induced cardiotoxicity, respectively^46,47^.

**Figure 5.**
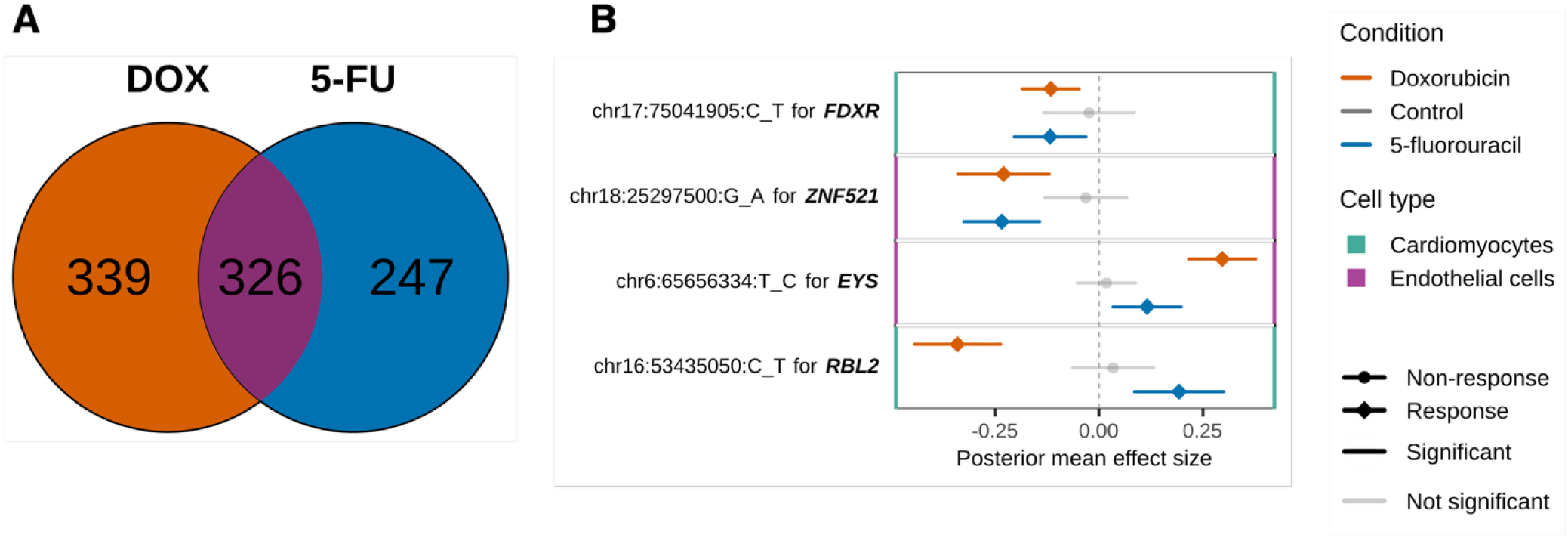
One third of response eGenes are shared between DOX and 5-FU, including known drug-induced cardiotoxicity risk genes. (**A)** 35.7% of all treatment-response eGenes are shared (purple) between the DOX (orange) and 5-FU (blue) treatments. (**B**) Forest plots showing effect size estimates for response corresponding to the eGenes *FDXR*, *ZNF521*, *EYS*, and *RBL2*. A diamond shape indicates response for a given cell-type-treatment condition and a circle indicates non-response. Effect size estimates for DOX, 5-FU, and control groups are given for the cell type indicated by the colored vertical borders on each panel. Per condition, significant effects (lfsr <0.1) are solid and nonsignificant effects are semi-transparent.

Intriguingly, a number of the eGenes identified in both DOX-treated and 5-FU-treated cells of the same type are differentially regulated. For example, a response eQTL for the tumor suppressor and transcriptional corepressor *RBL2* is associated with increased *RBL2* expression in 5-FU-treated cardiomyocytes; yet in cardiomyocytes treated with DOX, *RBL2* expression is downregulated. A previous study found that DOX-induced cardiotoxicity is exacerbated in *RBL2* knockout mice, which incur severe mitochondrial damage and cardiomyocyte apoptosis, resulting in reduced heart function and mass^48^. Although to our knowledge *RBL2* has not been linked to 5-FU-induced cardiotoxicity, the activity of this shared response eQTL suggests that the same variant with deleterious effects in the context of DOX may be cardioprotective in the context of 5-FU.

Our findings suggest that the transcriptional effects of DOX and 5-FU may converge on similar genes and pathways to perturb cardiovascular function, potentially through the same mechanisms that drive cardiovascular disease in the absence of drug-induced cardiotoxicity. Certain risk genes already linked to DOX-induced cardiotoxicity may warrant investigation as potential biomarkers for 5-FU.

### Drug-response eQTLs colocalize with GWAS signals relevant to cardiovascular disease

To formally assess the relevance of drug response eGenes to cardiovascular health, we used colocalization analysis^49^ to identify signals associated with both response eGene expression and cardiovascular disease (**Methods**). Specifically, we analyzed data from the VA Million Veteran Program study, which includes summary statistics from genome-wide association studies (**GWAS**) of myocardial infarction, congestive heart failure, ischemic heart disease, and other relevant traits^50^. We detected four signals representing credible colocalizations (posterior probability that both the GWAS and eQTL traits associate with the same signal [PP.H4] >0.5) between cardiovascular traits and response eGenes (**Figure 6**, **Supplementary Table 7**).

**Figure 6.**
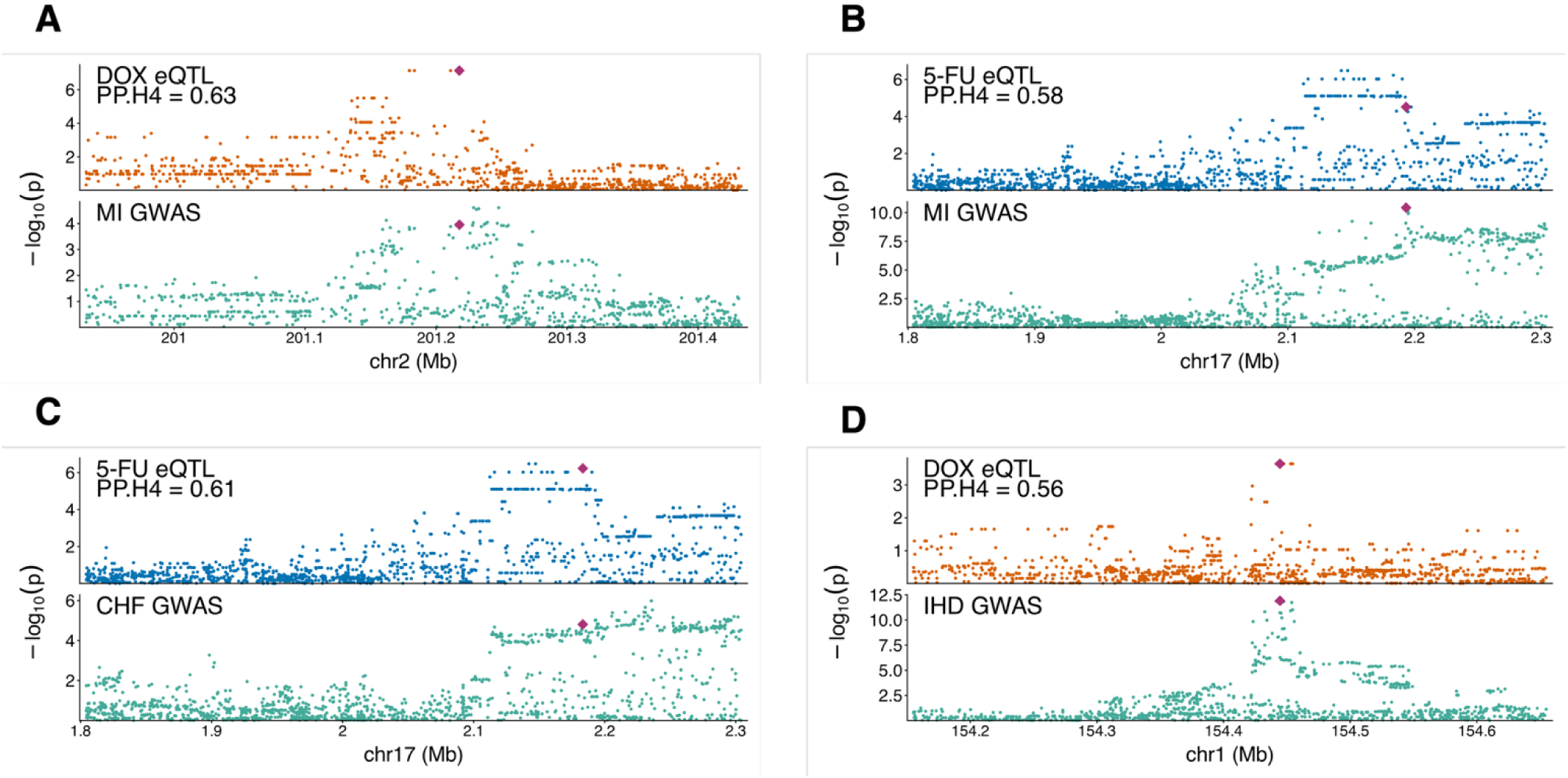
Colocalized signals associated with both cardiotoxicity-related diseases and eQTL-regulated gene expression changes. Myocardial infarction (MI) GWAS signals (green) colocalize with (**A**) rs13006529, a response eQTL for *CASP10* in DOX-treated cardiac fibroblasts (orange) and (**B**) rs6503321, a response eQTL for *HIC1* in 5-FU-treated proliferating fibroblasts (blue). (**C**) A GWAS signal for congestive heart failure (CHF; green) colocalizes with rs9908373, a response eQTL for *HIC1* in 5-FU-treated proliferating fibroblasts (blue). (**D**) A GWAS signal for ischemic heart disease (IHD; green) colocalizes with rs12730036, a response eQTL for *IL6R* in DOX-treated epithelial cells (orange). Lead candidate SNPs for each colocalization are indicated by a red diamond.

Two signals associated with myocardial infarction colocalize with the DOX response eGene *CASP10* (PP.H4 = 0.63) in cardiac fibroblasts (**Figure 6A**) and the 5-FU response eGene *HIC1* (PP.H4 = 0.61) in proliferating fibroblasts (**Figure 6B**). *CASP10*, involved in apoptosis, was initially considered to be pro-apoptotic^51^, though pro-survival functionality has been reported more recently^52^. *CASP10* polymorphisms have been linked to toxicity outcomes in platinum-based chemotherapy^53^, and in a recent study using primary cultures of adult ventricular heart tissue treated with DOX, *CASP10* was significantly upregulated relative to control cells^54^. In addition to myocardial infarction, the 5-FU-response eGene *HIC1* colocalizes (PP.H4 = 0.61) with an association for congestive heart failure in proliferating fibroblasts (**Figure 6C**). *HIC1* is a tumor suppressor and quiescence-associated factor marking mesenchymal progenitor cells^55^; its loss in stromal precursors was shown to drive arrhythmogenic cardiomyopathy in mice^56^ and was reported as a candidate gene involved in mitral valve prolapse^57^. Another locus associated with ischemic heart disease credibly colocalized with *IL6R* (PP.H4 = 0.56), a DOX response eGene in epithelial cells (**Figure 6D**). The IL6R cytokine receptor and its ligand (IL6) are core components of inflammation, and IL6-IL6R signaling is known to be proatherogenic^58^. Moreover, previous studies support IL6 as a potential prognostic biomarker for DOX-induced cardiotoxicity^59^.

Taken together, these data suggest that response eGenes identified in cardiac HDCs are relevant to both drug-induced cardiotoxicity and cardiovascular disease, underscoring the potential of response eQTLs to serve as genetic predictors of both drug-induced side effects and disease risk.

## DISCUSSION

A substantial body of work shows that context-specific eQTLs are more often related to differences between individuals in clinically relevant phenotypes, including disease risk, than baseline steady-state eQTLs^16,18,32,60–64^. Genetic variants that regulate genes expressed in the heart and vasculature are likely to modify inter-individual risk for drug-induced cardiotoxicity, particularly if the regulatory impact of the genetic variant is altered specifically in response to drug treatment. We used cardiac HDCs to explore how inter-individual differences in genetic background affect gene regulation in response to DOX and 5-FU, the top two causes of cardiotoxicity among chemotherapeutic drugs. DOX in particular is notorious for its adverse cardiac effects, which are often severe and affect a large proportion of patients. A growing body of *in vitro* work in iPSC-derived cardiomyocytes has helped to reveal genes and pathways that are responsive to DOX treatment *in vitro*^13–15^, many of which are likely to have *in vivo* relevance as biomarkers for predicting a patient’s risk for DOX-induced cardiotoxicity^46^. Our results support and expand upon previous studies of DOX by exploring its effects in a multicellular context.

In contrast to DOX, the molecular effects of 5-FU are surprisingly understudied for a drug that has been prescribed to treat cancer for over 60 years^2,3,11^. As the first study to characterize the transcriptomic effects of 5-FU across multiple cardiovascular cell types, our data provide a much-needed foundation that can be used to explore the drug’s putative mechanisms as well as the genetic basis of its effects. While many of the 5-FU response eGenes we identified have already been linked to drug-induced cardiotoxicity or disease, we also identified a number of genes whose putative cardiovascular effects are unknown, suggesting new avenues of investigation for future studies of 5-FU. We also found hundreds of eGenes that respond to both 5-FU and DOX, suggesting shared mechanisms that could be relevant to other drugs with cardiovascular side effects.

Our colocalization analyses demonstrate that the phenotypic responses expressed by cardiac HDCs are relevant to both drug-induced cardiotoxicity and long-term cardiovascular health. This result is consistent with the observation that patients treated with DOX and 5-FU are at greater risk for developing cardiovascular pathologies such as myocardial infarction and arrhythmia^5–7^. Although there is no GWAS that is well-powered enough for us to assess eQTL colocalization among loci directly associated with drug-induced cardiotoxicity, we did identify drug response eQTLs that colocalize with GWAS signals for myocardial infarction and congestive heart failure. Additionally, we found a signal for ischemic heart disease that colocalizes with the DOX-response eGene *IL6R* in epithelial cells. IL6R is the receptor for the cytokine IL6, which promotes inflammation by binding IL6R in its soluble form. While IL6 *trans* signaling can occur in any cell type via the ubiquitously expressed gp130 signal transducer, epithelial cells are one of the few that express membrane-bound IL6R. This gives epithelial cells an innate capacity for classic IL6 signaling, which supports the cytokine’s anti-inflammatory and regenerative functions. However, IL6R-expressing cells undergoing apoptosis also shed a soluble form of the receptor that then facilitates pathogenic inflammation in other cell types, including cardiovascular cells, through IL6 *trans* signaling^65^. Not only is IL6 signaling known to be proatherogenic^58^, but patient studies support IL6 as a potential prognostic biomarker for DOX-induced cardiotoxicity^59^. Furthermore, blockade of IL6R-IL6 binding using monoclonal antibodies (e.g., tocilizumab or ziltivekimab) stops both classic and *trans* signaling. This blockade is used to treat several diseases, including cardiovascular disorders^58,66^, and has been shown to attenuate immunotherapy-induced cardiotoxicity in mice^67^. Further investigation using the relevant monoclonal antibodies to block IL6R-IL6 binding may also provide insights for mitigating DOX-induced cardiotoxicity.

The use of cardiac HDCs allowed us to explore the effects of DOX and 5-FU across multiple cell types relevant to cardiotoxicity and cardiovascular disease, including cell types and states that have not been extensively studied in the context of chemotherapeutic drugs. Cardiac HDCs also enable heterotypic cell-to-cell signaling, which has been shown to better recapitulate *in vivo* physiology and support cell development compared to monolayer cultures. Heterotypic cell-to-cell signaling is not only pertinent to cardiac injury^68^, but it also provides morphogenic information and developmental cues that would be impractical to introduce exogenously. These features, together with the efficiency and simplicity of guided differentiation, make cardiac HDCs suitable for population-level genomic studies exploring the effects of environmental exposures on cardiovascular tissues. We expect that future studies of cardiac HDCs will reveal that other cardiotoxic drugs have wide-ranging and variable effects across cell types, similar to what we found for DOX and 5-FU. Even targeted drugs, which are theoretically designed to be safer than conventional chemotherapeutics, have been associated with cardiotoxicity; for example, tyrosine kinase inhibitors (e.g., imatinib mesylate) have damaging effects on both cardiomyocytes and endothelial cells^69,70^. Future studies of cardiac HDCs could determine whether the specificity of the transcriptional response varies across drug modalities, doses, and treatment regimes, which could help to inform the development of new therapeutics.

The cellular diversity present in cardiac HDCs is likely to hold considerable appeal for drug screening studies in general: out of the small fraction of drugs that pass clinical trials, nearly 10% are withdrawn from the market due to cardiotoxicity^71^, possibly due to effects that would be detectable by measuring responses across a broader range of cell types and states. Cardiac HDCs may prove to be particularly relevant in the current United States regulatory environment, where drugs are no longer required to be tested in animals before entering clinical trials^72^. Understanding who is at risk for drug-induced cardiotoxicity is critical, as adverse side effects can lead patients to prematurely terminate chemotherapy, reducing treatment efficacy and potentially causing death. The ongoing lack of consensus on biomarkers and genotype predictors underscores an urgent need for population-level studies of drug responses in diverse cell types and genetic backgrounds, which cardiac HDCs and other iPSC-based models are poised to fulfill.

## STUDY LIMITATIONS

In developing the cardiac HDC model, we intentionally prioritized efficiency and reproducibility over fidelity to adult cardiac tissues. Cardiac HDCs are therefore better-suited for eQTL mapping than existing cardiac organoids, but they do not recapitulate the structure or physiology of the heart. It has been repeatedly shown that eQTLs mapped in *in vitro* differentiated cells that are not fully matured recapitulate regulatory phenotypes seen in adult tissues^19,69,73,74^. In particular, studies of iPSC-derived cardiomyocytes treated with DOX have suggested that *in vitro* phenotypes are reasonable proxies for those observed in patients, at least on the molecular level^40^. Nevertheless, it is possible that the transcriptional responses observed in cardiac HDCs may differ from those observed in cancer patients.

A second limitation of our study is the use of a single concentration for each drug. These concentrations were selected based on previous reports of *in vitro* exposure as well as measurements of drug concentrations in patient plasma following treatment^74,75^. It is possible that a range of doses may have revealed additional effects beyond what we captured with a single concentration per drug. In designing our study, we weighed the benefits of multiple drugs versus multiple doses of a single drug. Given the degree of response eGene overlap between the two different treatments, we would expect even greater redundancy for multiple concentrations of the same drug. We therefore assess that, in retrospect, the choice of multiple drugs likely yielded the maximal results in terms of cardiotoxicity risk gene discovery. However, future studies may aim to explore drug effects empirically across a range of doses and this dataset will serve as a valuable benchmark, especially for *in vitro* studies of 5-FU.

## ACKNOWLEDGEMENTS

We thank all members of the Gilad lab for their support. This work was completed in part with resources provided by the University of Chicago’s Research Computing Center. We thank the University of Chicago Functional Genomics Core Facility for their assistance with sequencing the libraries. Figures were created using BioRender.com.

## AUTHOR CONTRIBUTIONS

Y.G. conceived the study and supervised the project. E.M. developed the guided differentiation method. E.M. designed the study with support from Y.G. and K.A.B. E.M. and O.L.A. performed all of the experiments and single-cell RNA-seq library preparation. J.B. provided support and valuable advice regarding cell culture. E.M. analyzed and interpreted the data with support from Y.G. and K.A.B. E.M. drafted the manuscript with review and editing provided by Y.G. and N.M.G.

## MATERIALS AND METHODS

### Samples

We used 49 iPSC lines (46 after quality control and filtering, 17 male and 29 female) from unrelated Yoruba individuals from Ibadan, Nigeria (YRI). These iPSC lines were reprogrammed from lymphoblastoid cell lines and were characterized and validated previously^20^. We confirmed cell line identities using genotype data generated by the 1000 Genomes Project^76^. WiCell’s Characterization Laboratory (Madison, WI) karyotyped all cell lines to assess genetic stability and confirm normal karyotypes.

### iPSC maintenance

We maintained iPSC lines using Matrigel Growth Factor Reduced (GFR) Basement Membrane Matrix (354230, Corning) with Essential 8 (E8) medium (A1517001, Thermo Fisher Scientific) and Penicillin-Streptomycin (17-602F, Lonza) in an incubator at 37°C and 5% CO2. At roughly 80% confluency (approximately every 3-5 days), we passaged cell cultures using a dissociation reagent (0.5 mM EDTA, 300 mM NaCl in PBS) and seeded iPSCs with 10 µM ROCK inhibitor Y-27632 (ab120129, Abcam).

### Guided differentiation and drug treatment

We formed and harvested cardiac HDCs across six batches, each consisting of 6-10 iPSC lines, balanced by sex and passage number. We performed guided differentiation of iPSCs by forming three-dimensional stem cell aggregates and applying temporal Wnt modulation. We dissolved the small molecules CHIR99021 (4423, Tocris) and IWP2 (3533, Tocris) in BloodStor 100 (100% DMSO; 410301, Biolife Solutions) at stock concentrations of 10 mM and 4 mM, respectively. We stored stock solutions at-20°C and remade them at least every three months. We formed iPSC aggregates using an AggreWell 800 24-well plate (34811, STEMCELL Technologies), coating each well with Anti-Adherence Rinsing Solution (07010, STEMCELL Technologies). On day 0, we dissociated iPSCs (using 0.5 mM EDTA, 300 mM NaCl in PBS) and seeded them onto the plate with E8 medium and 10 µM CEPT^77^ (i.e., chroman 1; 7163, Tocris), emricasan (510230, MedKoo Biosciences), polyamines (P8483, Sigma-Aldrich), and trans-ISRIB (5284, Tocris). To form aggregates we seeded the cells at a density of 2 million per well to target a 300 µm diameter^78^, the optimal size for inducing pluripotent stem cells towards the cardiac lineage^79^. We centrifuged the plate at 100 × g for 3 min to aggregate cells and maintained them in the plate for 24 h. After 24 h, we transferred aggregates to a suspension culture using ultra-low adherent plates (100-0083, STEMCELL Technologies) with 4 mL of medium per well.

On day 1 of guided differentiation, we placed cells in E8 medium for 48 h of ‘pre-differentiation’ to allow for recovery and reestablishment of cell-to-cell contact following formation (note that we maintained the cells in the initial E8 medium throughout the entire 48 h duration with no replenishment to allow a reduction in thermolabile FGF activity, thereby providing a transitional stage prior to initiating cell differentiation). On day 3, we switched the cell culture medium to ‘heart medium’ plus 6 µM CHIR99021. Heart medium consists of RPMI 1640 Medium, GlutaMAX Supplement, HEPES (72400047, Thermo Fisher Scientific), 2% v/v B-27 Supplement, minus insulin (A1895601, Thermo Fisher Scientific), and Penicillin-Streptomycin. After 24 h, on day 4, we changed the medium to base heart medium for 48 h.

On day 6, we changed the medium to heart medium plus 4 µM IWP2 for 48 h. From day 8 onward, cells were maintained in base heart medium, which we refreshed every 48 h (days 8, 10, 12, 14). Importantly, insulin is not introduced at any point during guided differentiation in order to maintain a culture environment permissive for nonmyocyte differentiation. On day 15, we performed a final base heart medium refresh wherein one of each of the three wells per cell line was treated with base heart medium containing either 1 µM DOX (Selleck Chemicals, S1208), 5 µM 5-FU (Selleck Chemicals, S1209), or a DMSO vehicle control (Sigma-Aldrich, D4550-100ML). All treatments were standardized to a 0.05% DMSO concentration.

### Aggregate dissociation and scRNA-seq

We collected cardiac HDCs on day 16 and dissociated them by exposing them to room temperature AccuMax (07921, STEMCELL Technologies) followed by incubation at 37°C for 10 min. Following incubation, we pipetted aggregates up-and-down for 30 sec with a p1000 wide-bore pipette tip (2079GPK, Thermo Fisher Scientific). We then incubated aggregates for an additional 5 min at 37°C. We repeated the pipetting every 5 min until aggregates were dissociated, at which point we added 5 mL of 4°C base heart medium and centrifuged cells for 100 × g for 3 min. We resuspended cells in 1 mL 4°C base heart medium and strained cells through a 40 µm cell strainer (H136800040, Bel-Art). We combined cells from each iPSC line in even proportions and centrifuged cells at 100 × g for 3 min. We combined cell suspensions from each treatment group at even proportions and centrifuged at 100 × g for 3 min. We then resuspended cells in 4°C base heart medium at a concentration of ∼1.6 million cells/mL. We collected cells from cardiac HDCs for scRNA-seq using the 10x Genomics Chromium Next GEM Single Cell 3’ HT Reagent Kits v3.1 (Dual Index; PN-1000348, 10x Genomics). For each batch, we used two lanes of a 10x chip, recovering ∼20,000 cells per lane (∼40,000 cells per batch). We sequenced the libraries using an Illumina NovaSeq X at the University of Chicago Functional Genomics Core Facility.

### scRNA-seq analysis

We captured transcriptional profiles from 49 iPSC lines, including 818,230 cells and 27.3 billion sequencing reads (33,401 reads-per-cell on average). We aligned samples to the human genome (GRCh38) using Cell Ranger (v7.0.0)^80^ and assigned cells to individuals using Vireo (v0.5.6)^81^. We removed two iPSC lines from batch 1 due to cross-contamination. Cell line NA19114 was mistakenly collected twice (in batches 1 and 5) due to mislabeling; we kept only the original collection for downstream analyses. We analyzed count data in R (v4.4.1) using Seurat (v5.3.0)^21^. We merged all Seurat objects per treatment group and removed cells classified as doublets or unassigned by Vireo; we also removed cells with <500 or>9,000 unique genes, as well as cells with >60,000 UMIs. We did not filter based on percentage of reads mapped to the mitochondrial genome; while mitochondrial percentage is often used as a proxy for dying cells, cardiomyocytes and hepatocytes naturally possess high levels of mitochondrial DNA and may be inadvertently removed using this technique^82^. After filtering, we retained 46 unique individuals within each treatment group with a total of 668,281 high-quality cells, including 217,276 control-treated cells, 208,092 DOX-treated cells, and 242,913 5-FU-treated cells.

We performed log-based normalization of gene expression for each treatment group independently. For the control data only, we performed scaling and regressed out cell cycle phase differences to minimize variability among proliferating cells while preserving phase differences between cycling and non-cycling cells, thereby maintaining distinctions between progenitor and terminally differentiated populations. We applied principal components analysis (**PCA**) and used the top 50 principal components (**PCs**) for UMAP embedding. We computed k-nearest neighbors and then performed unsupervised clustering at a resolution of 0.12, yielding 14 cell clusters for DE analysis using the Wilcoxon Rank Sum test.

We annotated cell clusters based on expression of key marker genes. Then, we transferred cell labels from the annotated control reference using the Seurat (v5.3.0) MapQuery function to annotate cells in the two drug treatment groups (DOX and 5-FU)^21^. First, for the DOX and 5-FU groups, we performed log-based normalization of gene expression and then carried out automated cell annotation using Seurat label transfer to classify query cells (from either DOX or 5-FU) based on reference data (manually annotated control cells). We first identified a set of transfer anchors between the reference and query datasets using the PCA dimensional reduction of our control cells. 50 dimensions from the control reference PCA were used to project the query dataset onto the structure of our control reference dataset and identify anchors (pairs of cells from each dataset acting as mutual nearest neighbors) and correct the query’s projected embeddings. We integrated using the precomputed anchors and projected each query dataset onto the UMAP structure of the control reference. Prediction scores were calculated for each annotated query cell. To validate the robustness of our automated cell annotation approach, we confirmed high cell prediction scores for both treatment datasets. Additionally, we evaluated gene expression of the classified drug treatment cell types to ensure cell-type-specific gene expression patterns matched the same key marker genes used for manual annotation of control cells.

### Differential expression and gene set enrichment analyses

We performed DE analysis between control-treated and either DOX-treated or 5-FU-treated cells within each cell type using dreamlet (v1.4.1)^83^ on pseudo-bulked scRNA-seq data and used the resulting gene-level statistics for pre-ranked Hallmark gene set enrichment analysis (**GSEA**). We performed pre-ranked Hallmark GSEA with fgsea (v1.32.4)^22^ on dreamlet-derived z.std statistics using gene sets containing 15-500 genes and 100,000 permutations. Specifically, we extracted raw RNA counts from the Seurat objects and converted these into SingleCellExperiment format (v1.28.1)^84^. We aggregated counts across cells from the same individual–treatment combination within cell types. We retained pseudobulk samples with ≥15 cells and cell types with ≥25 samples. We retained genes that expressed above a counts-per-million threshold corresponding to min.count = 2 in ≥25% of retained samples. We then used processAssays to normalize counts and estimate precision weights and fit linear mixed models with treatment, sex, and batch as fixed effects and cell line as a random effect.

### *cis*-eQTL mapping with MatrixEQTL

To prepare scRNA-seq data for *cis*-eQTL mapping, we identified cell types that are well-represented across individuals using a two-stage quality control strategy for each cell type × treatment condition. First, we required that every individual have ≥15 cells of a given type in a given treatment group. Second, we required that each cell type in a given treatment group be represented by ≥25 individuals to be retained. Twelve cell types failed to meet these criteria and were excluded from all three treatment groups: hematopoietic cells, neural crest cells, neural progenitor cells, and neuroectodermal cells. We retained ten cell types in each of the three treatment groups for downstream analysis: cardiac fibroblasts, activated fibroblasts, cardiomyocytes, epithelial cells, mesenchymal cells, epicardial cells, foregut endodermal cells, proliferating fibroblasts, endothelial cells, and hepatic endodermal cells. We retained only protein-coding genes with a median per-sample count of ≥ 1 and nonzero variance per cell type across cell type × treatment conditions. We extracted raw RNA counts from the Seurat objects; these data were then converted into SingleCellExperiment (v1.28.1)^84^ format to make the data more tractable for pseudo-bulking and downstream eQTL mapping. We used dreamlet (v1.4.1)^83^ to pseudo-bulk counts for each individual in each treatment group. We normalized counts per cell type using trimmed mean of M-values and converted these to logCPM with edgeR (v4.4.2)^85^. We then inverse-normal-transformed the normalized reads for each gene in each cell type × treatment condition.

We mapped *cis*-eQTLs for each cell type × treatment condition using MatrixEQTL (v2.3)^25^. We used whole-genome sequencing data from the 1000 Genomes Project to map *cis*-eQTLs^76^ after removing rare variants (MAF<0.05). We tested SNPs within a 250 kb mapping window of each gene’s TSS. We included sex as a covariate along with a select number of expression PCs to capture latent biological or technical variables. To calculate the optimal number of expression PCs to include as covariates, we iteratively mapped eQTLs using sex and an increasing number of expression PCs (0 through 20) as covariates to determine the number of PCs that yielded the maximum number of significant (Benjamini-Hochberg FDR < 0.1) eGenes for a given condition.

### *cis*-eQTL mapping with mash

We applied mash (v0.2.79)^26^ to our MatrixEQTL *cis*-eQTL results. We used all SNP-gene associations tested by MatrixEQTL (i.e., setting the p-value threshold to 1) and computed standard errors for each SNP-gene pair. We built a set of strong tests using the top SNP for each gene based on the lowest p-value across all cell type × treatment conditions. Mash facilitates joint analysis to share information across all 10 cell types and 3 treatments (a total of 30 different conditions), leveraging 12,927 independent SNP-gene pairs to assess a total of 387,810 effects. We then designated a random set using an unbiased (generally null) subset of all tests. We used the random set to produce canonical covariance matrices and used the expectation-maximization algorithm to learn the residual correlation structure that represented common sharing patterns across conditions (i.e., baseline noise). We used the strong set to learn the data-driven correlation structure, or the correlation structure of the true signal, by using the top 5 PCs to produce PCA-based covariance matrices; we also produced data-driven covariance matrices using flashier (v1.0.7)^86^ for empirical Bayes matrix factorization and constrained factors to non-negative values given the multi-tissue composition of our dataset. We used the PCA and empirical Bayes data-driven covariance matrices to initialize the extreme deconvolution algorithm and refine data-driven covariance matrix estimates. We fit the mash model to the random set to learn mixture weights on covariance matrices and scaling coefficients. Finally, we computed posterior summaries on the strong dataset by applying the fitted model to strong tests; this shrinks the strong test effect sizes towards the most credible estimates. We applied a threshold of lfsr < 0.1 for each SNP-gene pair to define standard *cis*-eQTLs as significant. We determined sharing of standard *cis*-eQTL effects across conditions based on if the standard *cis*-eQTL was significant in at least one of the two focal conditions and by magnitude (defined as sharing the same sign and within a factor of 2).

### Definition and identification of response eQTLs

We used a marginal eQTL test in place of an explicit genotype × treatment interaction model to identify response eQTLs. This approach leverages the posterior mean effect estimates output by mash, which employs a Bayesian framework to shrink the raw, unstable regression coefficients output by MatrixEQTL, retaining only the strongest signals^26^. The resulting posterior mean effects are effectively de-noised, allowing us to use the log-fold change in eQTL effects between conditions to distinguish between response and non-response eQTLs. We defined response eQTLs by comparing eQTL effects in control-treated vs. DOX-treated (or control-treated vs. 5-FU-treated) cells, one cell type at a time. We required that response eQTLs be: 1) present in both the control group and the focal treatment group, 2) significant (lsfr <0.1) in at least one of the two treatment groups (i.e., control vs the focal drug), and 3) have a fold change of effect size magnitude >4 (this is the threshold at which we observed response and non-response eQTLs to be distributed as expected throughout the TSS window^32^, as shown in **Supplementary Figures 7-8**). Also, if the eQTL was significant in both groups and had differently signed effect sizes, we counted it as a response eQTL. If an eQTL was present in both treatment groups and significant in at least one, but otherwise did not meet the magnitude threshold or sign requirements, we designated it as a non-response eQTL. We then defined eGenes as response or non-response eGenes according to the type of eQTL they associated with.

### Functional annotation of *cis*-eQTLs

We performed functional annotation of DOX and 5-FU-response and non-response eQTLs to determine their proximity to TSS and enrichment of regulatory elements. We mapped distance to TSS of response and non-response eQTLs within 250 kb upstream and downstream of the TSS across all treatment groups.

Within each treatment group, an eQTL may be classified as responsive in one cell type and non-responsive in another. However, for these analyses, we wanted to avoid counting eQTLs multiple times (in order to keep the responsive and non-responsive groups mutually exclusive). We therefore removed any non-response eQTLs that were present in <u>both</u> the control group and the focal treatment group (DOX and 5-FU groups were separately compared to the control group). Also, if an eQTL was responsive in <u>any</u> DOX-treated cell type (or, separately, in any 5-FU-treated cell type), we labeled it as a response eQTL. If a response eQTL was identified as a non-response eQTL in another cell type, it was only counted once – as a response eQTL.

Based on strand-aware GRCh38 gene annotation, we used the genomic coordinates to calculate eQTL distance relative to the corresponding eGene TSS; we then used these distance values to plot histograms. We calculated kernel density curves for each histogram and rescaled them to indicate distribution shape. For regulatory element enrichment, we leveraged ENCODE candidate *cis*-regulatory elements (**cCREs**)^87^. We measured enrichment of response and non-response eQTL SNPs relative to all other SNPs tested in the corresponding MatrixEQTL *cis*-eQTL analysis (provided in **Supplementary File 1**) for overlap with promoter-like signatures, proximal enhancer-like signatures, and distal enhancer-like signatures. Additionally, we assessed enhancer-like signatures overall (i.e., the union of proximal and distal enhancer-like signatures). We collapsed significant SNP-gene pairs generated from the mash analysis into unique eQTLs to perform enrichment analysis at the SNP level (meaning we did not count the same SNP-gene pair twice). For each treatment group, we performed two-sided Fisher’s exact tests separately to compare response and non-response eQTLs, comparing the proportion of each eQTL type that overlapped each cCRE category with its proportion of overlap among all other tested SNPs and calculated odds ratios, 95% confidence intervals, and p-values.

### Colocalization of GWAS signals and eQTLs

We performed colocalization analysis of standard eQTLs with case-control summary statistics for eight GWAS traits (odds ratios)^50^. Three traits showed plausible colocalization of GWAS signals with drug-response eGenes: myocardial infarction (Study: GCST90479551), congestive heart failure (Study: GCST90480181), ischemic heart disease (Study: GCST90480133). Five traits did not demonstrate colocalization of GWAS signals with any drug-response eGenes: atrial fibrillation and flutter (Study: GCST90480170), endocarditis (Study: GCST90480146), hypertension (Study: GCST90479553), pericarditis (Study: GCST90480144), and deep vein thrombosis (Study: GCST90480220). For each condition from our eQTL data, we selected those eGenes with a significant *cis-*eQTL (as defined above) and then extracted each eGene’s corresponding SNPs which were tested for association in MatrixEQTL (i.e., SNPs within 250 kb of the eGene’s TSS), along with each SNP’s MatrixEQTL statistics, including effect size beta, standard error, p-value, rsID, and effect size variance. We performed colocalization using the R package coloc (v5.2.3)^49^, taking an approximate Bayes factor (**ABF**) colocalization approach and assuming a single causal variant with default prior values. Coloc^49^ analyzes one genomic region at a time (in this case, 500 kb) and does not assume similar patterns of linkage disequilibrium between the GWAS and eQTL studies. The ABF colocalization method computes five posterior probabilities (**PP**) per region tested: no association in the region for either trait (PP.H0.ABF), only the GWAS trait has an association (PP.H1.ABF), only the eQTL trait has an association (PP.H2.ABF), both the GWAS and eQTL traits are associated with the region but with different causal variants (PP.H3.ABF), or both the GWAS and eQTL traits are associated with the same single causal variant (PP.H4.ABF). This allows for a wider range of interpretations with regard to how the GWAS and eQTL data relate to one another. If limited power is an issue, this is indicated by high H0, H1, or H2 values. We only considered those regions with a colocalization PP.H4.ABF of >0.5 to be colocalized.

